# Population genomic analysis of an emerging pathogen *Lonsdalea quercina* affecting various species of oaks in western North America

**DOI:** 10.1101/2023.01.20.524998

**Authors:** Olga Kozhar, Rachael A. Sitz, Reed Woyda, Lillian Legg, Jorge R. Ibarra Caballero, Ian S. Pearse, Zaid Abdo, Jane E. Stewart

**Affiliations:** Department of Agricultural Biology, Colorado State University, Fort Collins, CO, USA; Davey Resource Group, Inc., Urban & Community Forestry Services, Atascadero, CA, U.S.A.; Program of Cell and Molecular Biology, Colorado State University, Fort Collins, Colorado, USA; Department of Microbiology, Immunology and Pathology, College of Veterinary Medicine and Biomedical Sciences, Colorado State University, Fort Collins, CO, U.S.A.; U.S. Geological Survey, Fort Collins Science Center, Ft Collins, CO, USA

## Abstract

Previously unrecognized diseases continue to threaten the health of forest ecosystems globally. Understanding processes leading to disease emergence is important for effective disease management and prevention of future epidemics. Utilizing whole genome sequencing, we studied the phylogenetic relationship and within diversity of two populations of the bacterial oak pathogen *Lonsdalea quercina* from western North America (Colorado and California) and compared these populations to other *Lonsdalea* species found worldwide. Phylogenetic analysis separated Colorado and California populations into two well supported clades within the genus *Lonsdalea*, with an average nucleotide identity between them near species boundaries (95.31%) for bacteria, suggesting long isolation. Populations comprise distinct patterns in genetic structure and distribution. Genotypes collected from different host species and habitats were randomly distributed within the California cluster, while most Colorado isolates from introduced planted trees were distinct from isolates collected from a natural stand of CO native *Q. gambelii*, indicating the presence of cryptic population structure. The distribution of clones in California varied, while Colorado clones were always collected from neighboring trees. Despite its recent emergence, the Colorado population had higher nucleotide diversity, possibly due to migrants moving with nursery stock. Overall results suggest independent pathogen emergence in two states likely driven by changes in host-microbe interactions due to ecosystems conditions changes. To our knowledge, this is the first study on *L. quercina* population structure. Further studies are warranted to understand evolutionary relationships among *L. quercina* populations from different areas, including the native habitat of red oak in northeastern USA.

**Importance:** Bacterial pathogens from genus *Lonsdalea* severely affect oak forest ecosystems worldwide. In Colorado, USA, *L. quercina* is one of the causal agents of drippy blight disease on introduced red oak trees. Prior to discovery of drippy blight in Colorado, *L. quercina* was reported on oak trees in California, causing drippy nut on acorns of native oaks. Due to its recent emergence in Colorado, the origin and movement of *L. quercina* are unknown. In this study we investigated evolutionary relationships within genus *Lonsdalea* worldwide and *L. quercina* population structure in western USA. Our results demonstrate that *L. quercina* Colorado and California populations comprise distinct patterns of genetic structure and distribution, suggesting that accidental pathogen introduction from California to Colorado is unlikely. Higher nucleotide diversity in a recently emerged Colorado population suggests the bacterial strains might be migrants that initially moved with nursery stock from other areas in the last century. For example, Colorado strains of *L. quercina* may have moved from native stands of red oaks in the northeastern or southern USA. Curiously, however, this disease is not known in native red oak in the northeastern USA. Initial causes of recent disease emergence are likely driven by environmental/ecosystem changes since isolates for this study were collected from established mature trees. Results presented here give a better understanding of population biology of the bacterial oak pathogen and provide a framework for investigation of evolutionary relationships among pathogen populations from different areas.

## Introduction

Global forest ecosystems have been increasingly affected by a variety of disturbances, including altered climatic conditions and increased attack of key tree species by pests and pathogens. While the introduction of alien pathogen species to new geographic areas has been considered one of the main drivers of emerging forest diseases during the last century, other factors such as changes in environmental conditions, new host-vector associations (between introduced insect vectors and native tree pathogens or vice versa), cryptic disease agents (e.g., pathogens with a very long latency period or endophytes changing their behavior to pathogenic), hypervirulent strains of known pathogen species, and/or newly emerging species of unknown origin are also recognized as key factors leading to disease emergence in forest ecosystems around the globe (1–4). The complex nature of many forest diseases makes them difficult to manage and study. However, knowing underlying causes of disease emergence is crucial to facilitate effective disease management and preventing epidemic outbreaks that can have catastrophic consequences for ecosystems health.

The genus *Quercus* is one of the most important groups of trees in many regions of the Northern Hemisphere (5), with nearly 500 species that have been characterized globally. In addition, oak forests are characterized by high species diversity and great soil fertility(6). In North America oak trees compose a significant part of many forest ecosystems. For example, in the eastern USA, oak forest types represent roughly half of all forest land (7). However, the biodiversity of oak species varies within the continent, with roughly 90 species described in eastern North America, 20 species in California (5), and only one native species in Colorado. In North America, oak species are also often planted as shade trees in urban environments because of the ability to withstand harsh urban environments (8). This has led to the movement of oak species to new geographic areas. For example, northern red oak (*Quercus rubra*) that is native to eastern North America, has become a popular shade tree in Colorado, USA. These planted trees may experience new pathogens and/or pathogens may also act as bellwethers of the effects of climate change because they experience environmental conditions outside of their native range.

Oak decline diseases and complex syndromes, caused by combined effects of biotic and abiotic factors, have been reshaping landscapes of deciduous forests in temperate zones around the world (9). In the United Kingdom, acute oak decline has been associated with galleries of the two spotted oak borer, *Agrillus biguttatus*, and several bacterial species (most commonly with *Brenneria gudwini, Glibsiella quercinecans*, and *Rahnella victoriana*, as well as *Pseudomonas daroniae* and *P. dryadis*, and *Lonsdalea britannica)* (4, 10, 11). *Brenneria gudwini* and *G. quercinecans*, and *Brenneria* spp and *R. victoriana* are also associated with oak decline in Spain and Iran, respectively (10, 12).

In the early 2000s a significant dieback of northern red oak, pin oak, and Shumard oak of unknown origin appeared in Colorado and was described as drippy blight disease (13).

Recently emerged drippy blight on planted oaks causes abundant ooze on symptomatic tissue and has led to significant increases in tree mortality and removals in public properties in some areas of Colorado (13). Based on phenotypic, genetic analyses and pathogenicity tests it was confirmed that the disease was caused by the bacterial pathogen *Lonsdalea quercina* in association with scale kermes insect, *Allokermes galliformis*. Due to the development of abundant ooze on symptomatic tissue, it was called drippy blight disease. The first record of *L. quercina* was published in 1967 (14) reporting drippy nut disease on native oak trees in California, USA, that infects acorns and acorn caps of native oaks (*Q, agrifolia* and *Q. wislizeni*) probably using wounds made by wasps, weevils, and other common oak pests to enter the host tissue.

Besides *L. quercina*, three other species of *Lonsdalea*, all tree pathogens, have been described around the world: *L. iberica* causing bark canker and drippy nut disease on native oaks in Spain (15, 16), *L. britannica* associated with oak decline in Britain (15, 17), and *L. populi* causing canker disease of *Populus* in Spain and China (18, 19).

While the causal agents of drippy blight have been identified, the causes leading to its (and other *Lonsdalea* spp. oak pathogens) recent emergence remain unknown. To our knowledge, little is known about *L. quercina* population biology and relationship between the bacterial populations from different areas. Since the only two places in North America with published records of *L. quercina* causing disease on oak trees are California and Colorado, we studied the relationship between populations of *L. quercina* from oaks in California and Colorado. The objectives of the study were to determine the phylogenetic relationship between *L. quercina* populations from California and Colorado and other *Lonsdalea* species found worldwide and investigate population structure of *L. quercina* in these two locations. Since *L. quercina* was first documented in California, we also assessed whether the Colorado population originated from California, as well as discuss potential processes leading to the disease emergence in Colorado.

## RESULTS

### Pangenome and clonality

Our genome assembly and annotation utilized the Prokka and Roary analyses. The pangenome of combined California (CA; n = 52) and Colorado (CO; n = 31) populations consisted of 8,624 genes, 2,370 of which comprised the core genome (total of 2,460,085 bp) (Fig. 1). The core genomes of each population separately also contained a population specific set of genes that included 245 core genes for CA isolates and 372 core genes in CO (Fig. 1). Among the total 83 isolates sequenced there were 65 unique genotypes (46 in CA and 19 in CO; genotypic diversity – 88.46% in CA and 61.29% in CO) detected (Table 1, Table S1).

**Table 1.**
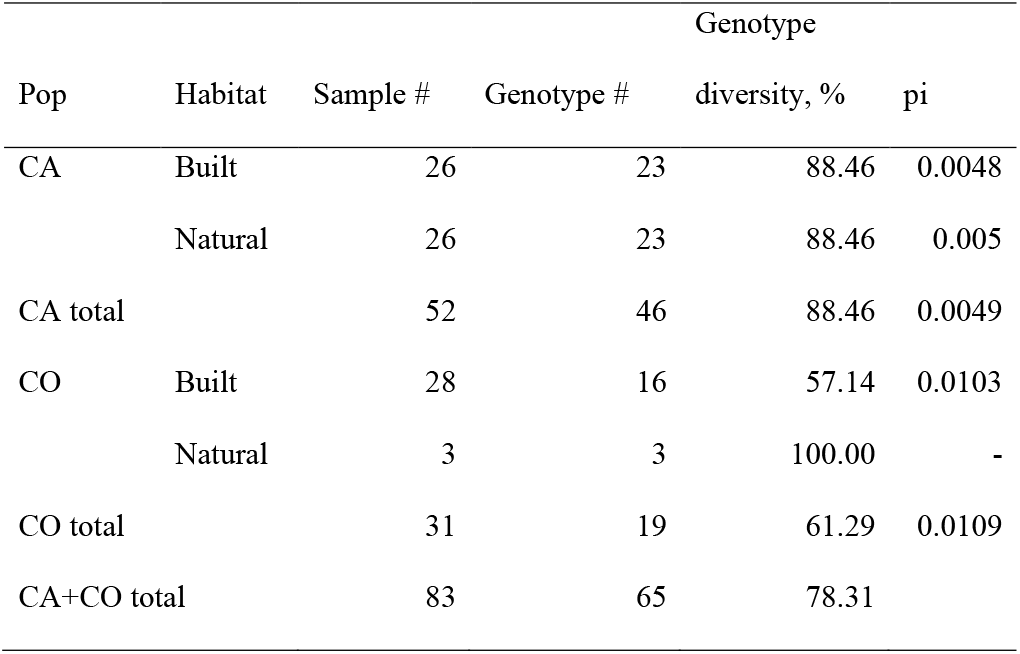
Diversity of *Lonsdalea quercina* populations in California (CA) and Colorado (CO), USA

**Figure 1.**
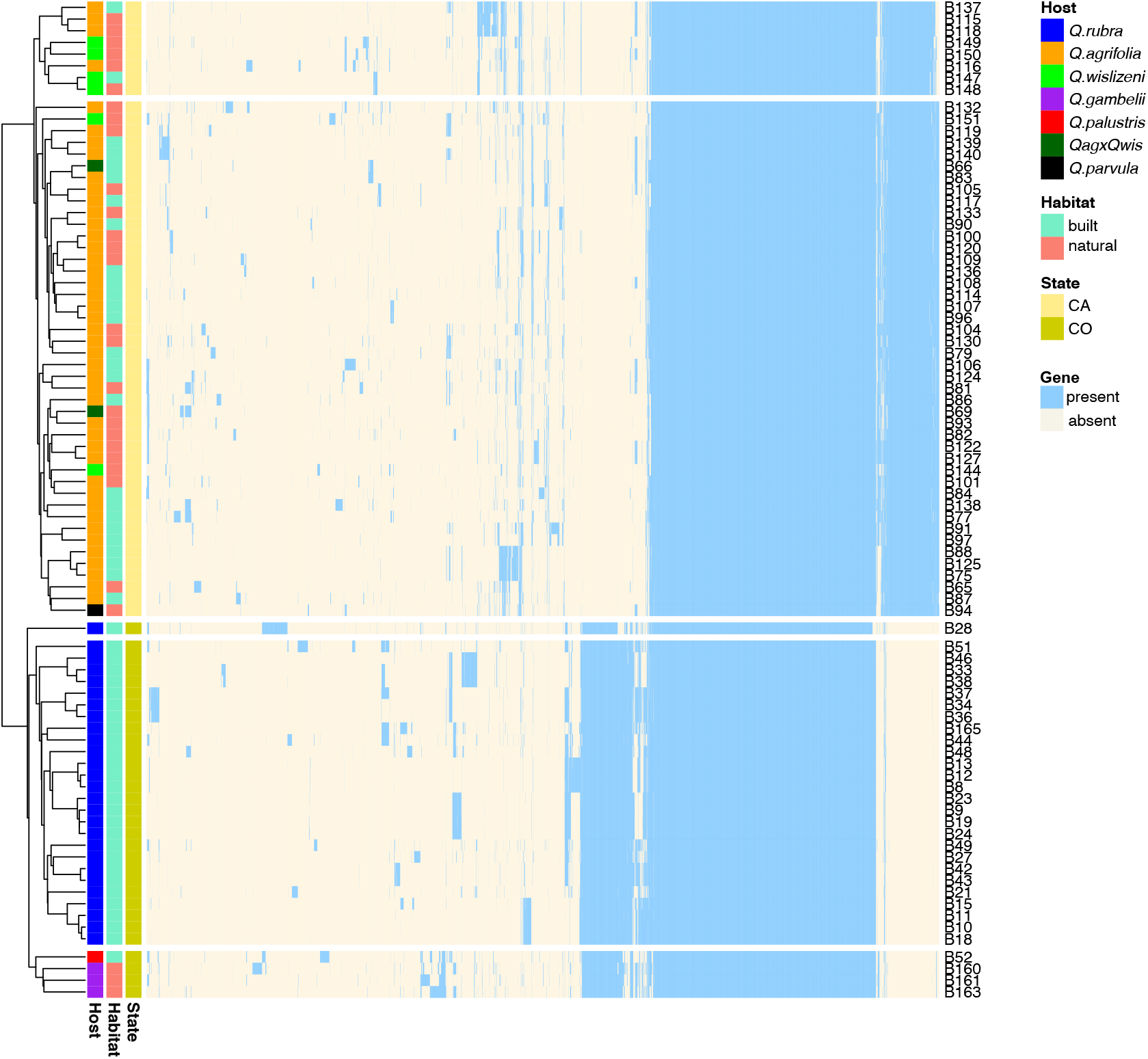
Gene presence-absence matrix from pangenome analysis of 83 isolates of *Lonsdalea quercina* populations collected in California (CA) and Colorado (CO), USA. Each column indicates a gene (blue – present, yellow – absent), each row represents a gene profile of each isolate. Host species, habitat, and state from which each isolate was collected are color coded according to the legend. Pangenome analysis was conducted with Roary v.3.13.0.

### Phylogenetic analyses and species boundaries

Phylogenetic analysis with other members of *Enterobacteriacea* family separated CA and CO populations into two distinct clades grouping them with two other previously published genomes of *Lonsdalea quercina*: members of the CA population were grouped with type strain ATTC 29281 previously collected in CA, and CO population genomes grouped with NCCB 100490 previously collected in CO (Fig. 2, Fig. S1). The phylogenetic analysis indicated a clear distinction between CA and CO populations of *L. quercina*, supporting grouping based on core gene presence-absence matrix (Fig. 1).

**Figure 2.**
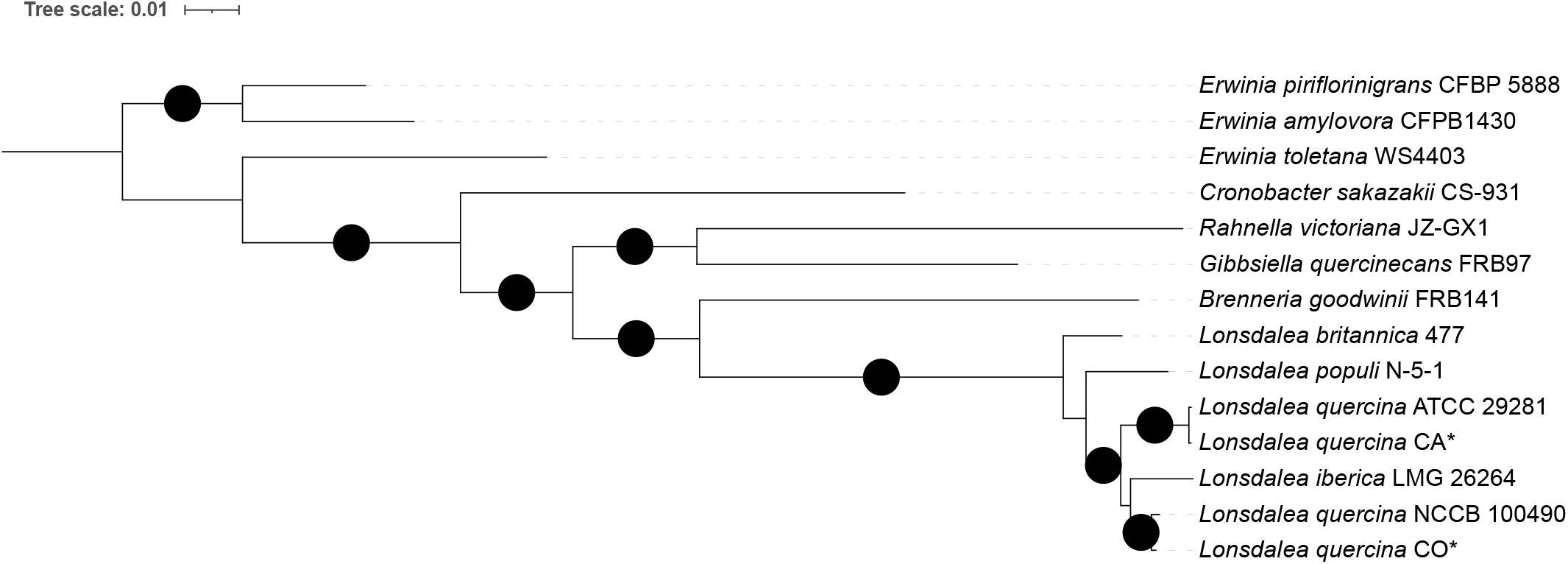
Maximum likelihood phylogenetic tree, based on 20 core genes shared among all individuals in the dataset, showing position of *Lonsdalea quercina* populations sampled in California (CA) and Colorado (CO) within genus *Lonsdalea* of family *Enterobacteriacea*. Branches with >=95% likelihood support are indicated with black circles. Branch support was calculated with 1000 ultrafast bootstrap replicates. *Erwinia piriflorinigrans* and *Erwinia amylovora* were used as root. \**Lonsdalea quercina* CA and *Lonsdalea quercina* CO are consensuses of multi-FASTA alignments of core genes of CA and CO populations, respectively, produced by Roary v.3.13.0.

Average nucleotide identity (ANI) above 95-96% is used to define bacterial species boundaries (20). Pairwise comparisons of ANI between consensus alignments of the core genomes of either CA or CO populations and one representative each of other *Lonsdalea* spp. were below the species boundaries threshold (between 89.08% - 93.75%), indicating a clear species boundary between *L. quercina* from other *Lonsdalea* species (Table 2). However, the average ANI between CA and CO was 95.18% (range 94.83 – 95.36) which is right on the threshold of ANI bacterial species boundaries. The CO population had a slightly lower within population average nucleotide identity (ANI) compared to CA population (98.98% for CO and 99.51% for CA, respectively). The lowest ANI between CA and CO populations was observed when a CO genome B28 was compared to any CA genome (average ANI between B28 genome and all CA genomes = 94.87%, range 94.83 – 95.01). ANI between B28 and CO population was lower than average within CO population ANI but was still within species boundaries (average ANI between B28 and the rest of CO genomes = 97.97%). When B28 genome was excluded from the analysis, the minimum ANI between any CA and CO genomes was 95.06%. These results suggest that CA and CO populations have being isolated from each other for a long time but may still be undergoing the process of allopatric speciation.

**Table 2.**
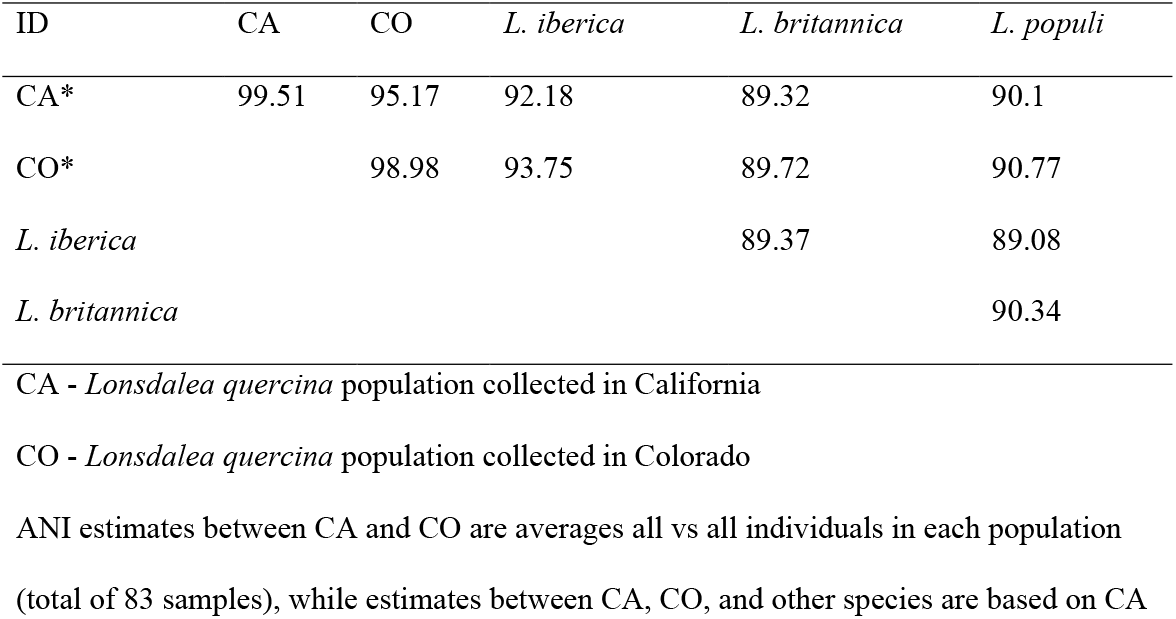
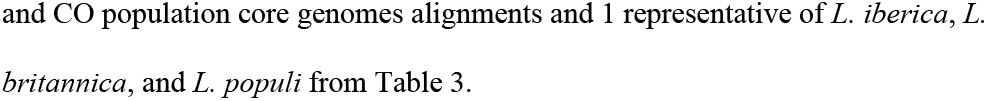
Estimates of Average Nucleotide Identity (ANI) within *Lonsdalea* genus and CO population core genomes alignments and 1 representative of *L. iberica, L. britannica*, and *L. populi* from Table 3.

| ID | CA | CO | *L. iberica* | *L. britannica* | *L. populi* |
| --- | --- | --- | --- | --- | --- |
| CA* | 99.51 | 95.17 | 92.18 | 89.32 | 90.1 |
| CO* |  | 98.98 | 93.75 | 89.72 | 90.77 |
| *L. iberica* |  |  |  | 89.37 | 89.08 |
| *L. britannica* |  |  |  |  | 90.34 |

CA - *Lonsdalea quercina* population collected in California
CO - *Lonsdalea quercina* population collected in Colorado
ANI estimates between CA and CO are averages all vs all individuals in each population
(total of 83 samples), while estimates between CA, CO, and other species are based on CA

### Population structure and distribution

Based on the gene presence-absence matrix of core and accessory genes CA and CO populations formed two separate clusters (Fig. 1). Both populations contained isolates from trees grown in natural (undeveloped landscapes such as forests, parks, nature preserves, etc.) or built (developed landscapes such as residential neighborhoods of cities, parking lots, etc.) areas (see materials and methods). CA population isolates from either natural or built areas were collected from different host species and these were randomly distributed within the CA cluster (Fig. 1, Table S1). In the CO population all samples, except 1, collected from planted trees grown in built areas formed a separate group, while three samples collected from native *Q. gambelii* in natural area were grouped together with 1 sample from a planted tree of a different species (*Q. palustris*). Due to the small sample size from natural area, it remains unclear whether bacterial populations are structured by habitat or host type in CO.

To further study population structure of CA and CO populations, we used an unrooted maximum likelihood phylogenetic tree (Fig. 3a). Consistent with the groupings in the gene presence-absence matrix (Fig. 1), CA and CO populations were separated into two well-supported clades, with isolates from different habitats and host species being randomly distributed within CA clade. Within CO clade, in turn, three isolates from native *Q. gambelli* (B160, B161, and B163) and 1 isolate from a planted *Q. palustris* (B52) formed a separate from planted *Q. rubra* isolates well supported cluster. Phylogenetic analyses also supported the gene presence-absence matrix grouping results for CO isolate B28 collected from a planted *Q. rubra* in a built area as genetically distinct from the rest of CO population.

**Figure 3.**
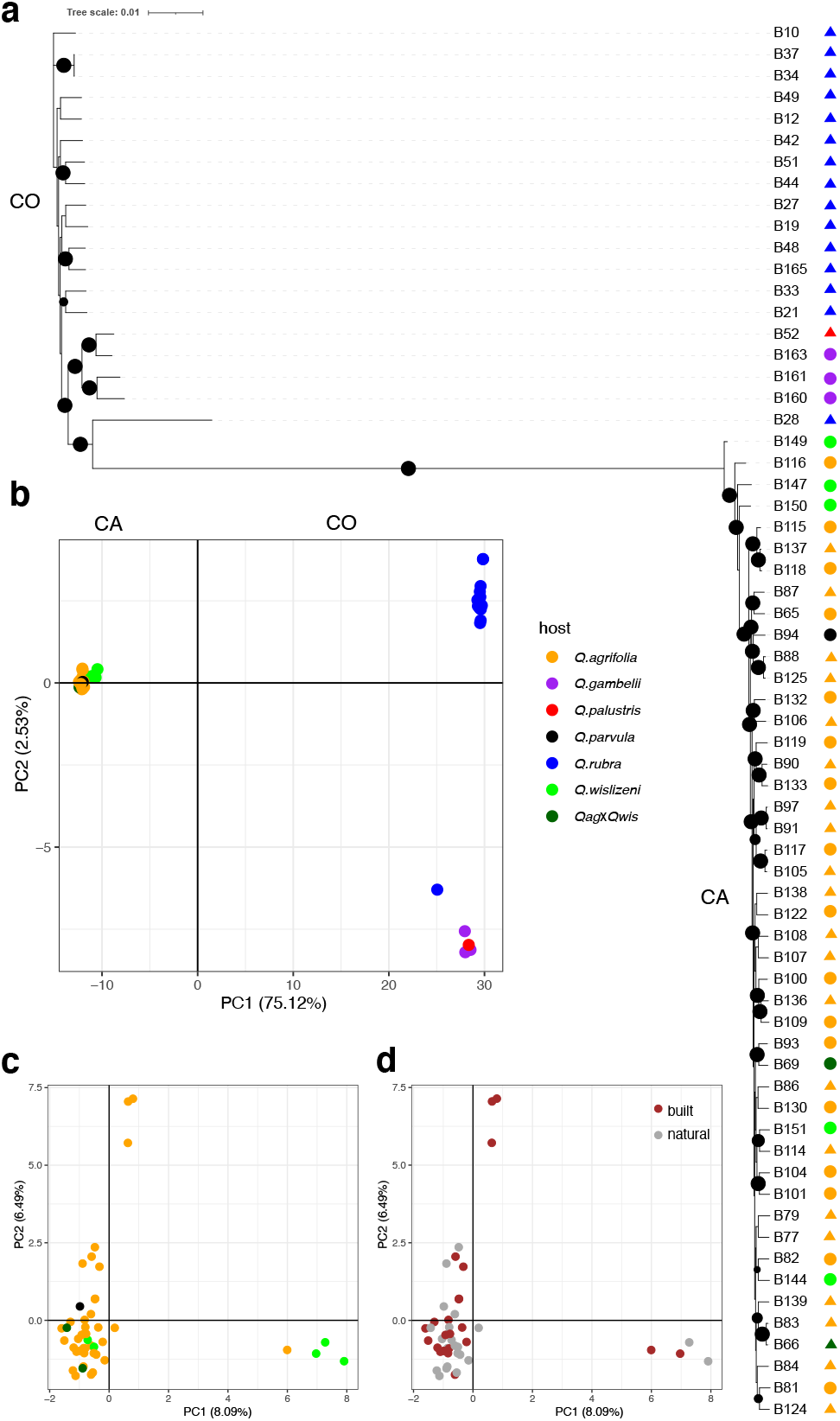
Population structure of *Lonsdalea quercina* sampled in California (CA) and Colorado (CO). **a**. Maximum likelihood phylogenetic tree showing distinct clades of CA and CO genotypes. Branches with >=95% likelihood support are indicated with black circles. Branch support was calculated with 1000 ultrafast bootstrap replicates. Circles and triangles next to phylogenetic tree indicate samples from natural and build habitats, respectively. *Q*.*agrifolia, Q. parvula, Q. wislizeni, and Q*.*agrifolia* x *Q. wislizeni* (QagxQwis) were sampled in CA, while *Q. rubra, Q. gambelii*, and *Q. palustris* - in CO. **b**. Principal component analysis (PCA) representing distinct genetic groups of CA and CO populations. **c, d**. PCA of CA population only representing lack of population structure either by host species (c) or habitat (d) within the state. Each dot indicates one individual. Percentages in parenthesis on PCA plots indicate the variance explained by each principal component. Different colors in **a, b**, and **c** represent isolates from different host species indicated in “host” legend. Different colors in **d** represent isolates from different habitats (“built”, “natural”).

Principal component analysis (PCA) based on unlinked core SNPs supported the results of phylogenetic analysis placing CA and CO populations into two distinct groups. CA isolates collected from various species of native oak and different habitats grouped closely together while more diversity was observed in CO population (Fig. 3b). In agreement with the groupings in the gene presence-absence matrix and maximum likelihood phylogenetic tree (Figs. 1 and 3a), three CO isolates collected from three plants of a native shrub *Q. gambelli* and 1 isolate from planted *Q. palustris* clustered together on the PCA plot, while rest of CO isolates (except B28) collected from planted *Q. rubra* grouped on the opposite side, further suggesting the presence of population structure within CO. Within CA population, no population structures either by host species of habitat was observed (Fig. 3c-d).

In CA, clones of 6 genotypes varied in their geographical distribution. Some representatives of the same genotype were collected from neighboring trees (Fig. 4a). For example, isolates of the B122 genotype were collected from trees located 4km from each other (both in natural areas - one in Mare Island Shoreline Heritage Preserve and another in Crockett Hills Regional Park) in San Pablo Bay area. Similarly, two isolates of the B139 genotype were collected from trees 80m apart in a built area on UC Davis campus. In contrast, representatives of another 4 genotypes were collected from geographically distant areas. Of these, two isolates each representing genotypes B125 and B107 were collected from trees in built areas located 35 and 83km apart, respectively (trees grown in parking lots or in highly developed city neighborhoods). Two isolates of genotype B100 were collected from trees in natural areas located 197km from each other (one in Mare Island Shoreline Heritage Preserve in San Pablo Bay area and another in Hastings Natural History Reservation in Carmel Valley). And finally, two isolates of genotype B147 were collected in oak savannah foothills on Rough and Ready highway outside of Yuba City – one alongside the road free of any development (categorized as natural area), and another 13km down the road next to a private house (categorized as built area) (Fig.4a). The CO population had higher clonality, but all representatives of the same genotype were always collected from trees in a close proximity to each other (e.g., isolates of either B8, B9, or B10 genotypes were collected around campus of CU Boulder) (Fig. 4b). Per site nucleotide diversity (pi) of CO was twice that of the CA population (0.0109 in CO and 0.0049 in CA, Table 1). Per site nucleotide diversity between isolates from CA built and natural areas did not differ (0.0048 and 0.0050, respectively).

**Figure 4.**
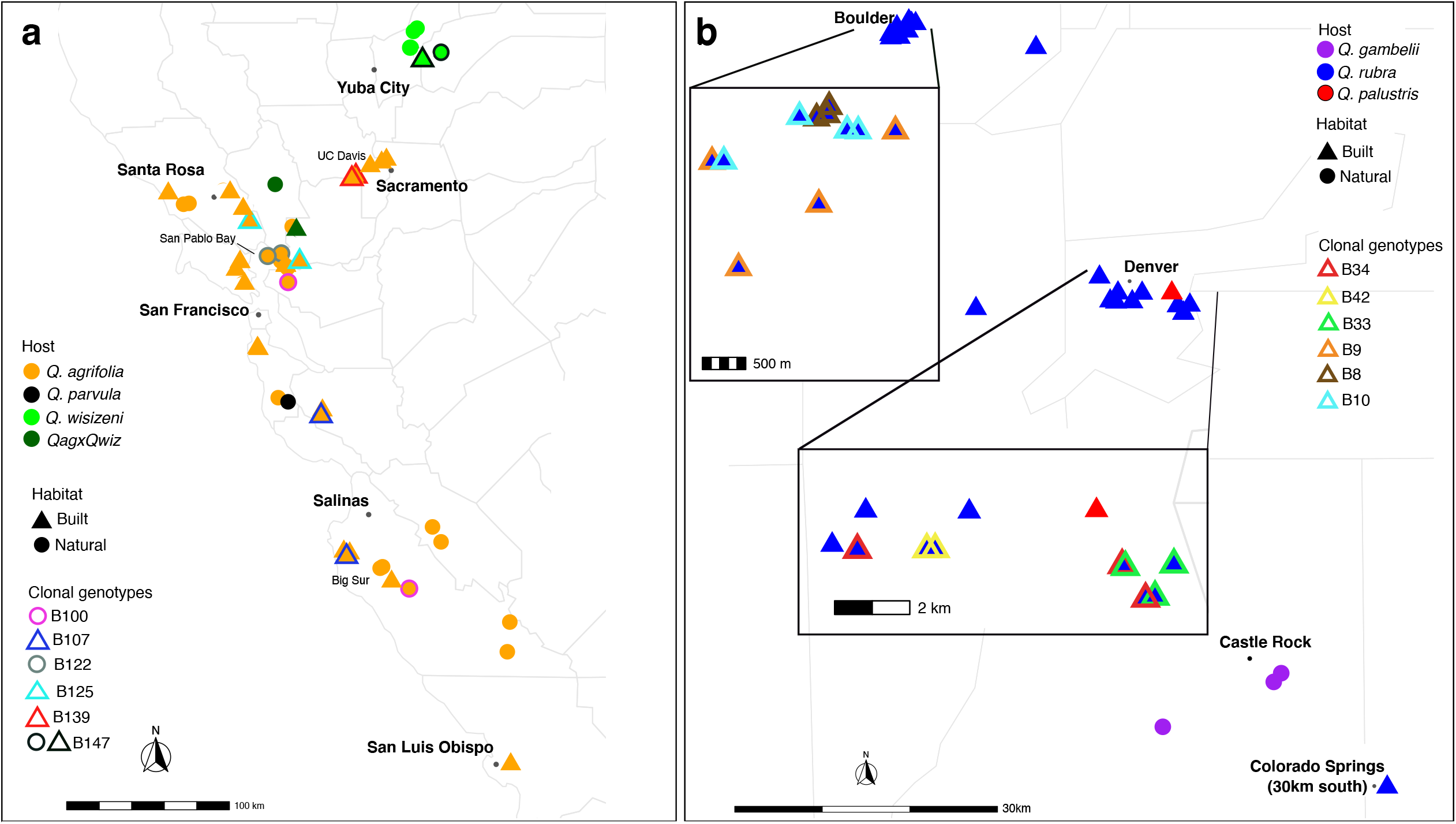
Distribution of *Lonsdalea quercina* populations in California **(a)** and Colorado **(b)**, USA. Every symbol on the map represents one isolate. Different colors represent different host species, and figures - habitats (see legends). Clonal genotypes are highlighted with different colors according to the legend “Clonal genotypes”. Unhighlighted samples are unique genotypes. Note different scale of each map. On Colorado map (b) Boulder and Denver areas are zoomed in to show clonal distribution within cities areas. One isolate from Colorado Springs (B165) is indicated in the right bottom corner (its actual location is 30km south).

### Recombination

The phylogenetic clustering of CA and CO populations was well supported with high bootstrap support, but due to unexpectedly higher nucleotide diversity in the recently emerged CO population compared to the CA population, we explored the possibility of recombination. The Phi test, based on 172,269 informative sites, revealed statistically significant evidence of recombination (p < 0.0001). Split decomposition analysis (21) suggested this was due to recombination among 3 isolates from native *Q. gambelli*, 1 isolate from *Q. palustris*, and 1 isolate (B28) from planted *Q. rubra* in the CO population, and two recombination events between isolates B48 and B165, B44 and B51, as well as several recombination events within CA populations (Fig. S2a-c). However, no recombination was detected between CA and CO populations. Relative rates of recombination to mutation were the same in CA and CO (R/θ = 0.149 for each), suggesting that genetic changes due to mutation are 7 times more likely than due to recombination in these populations. A ClonalFrameML phylogeny with branch length corrected to account for recombination showed longer branches in CO population compared to CA population, supporting results of higher nucleotide diversity in CO population (Fig. S3).

## Discussion

An increasing number of tree diseases cause significant, long term, threats to wild and human mediated ecosystems. Clarifying processes leading to the emergence of new diseases is a critical step towards their effective management. In this paper, we aim towards characterizing possible causes of emergence of drippy blight disease caused by *Lonsdalea quercina*, a recently emerged pathogen of oak species in association with kermes scale *Allokermis galliformis*, in Colorado, USA. Using population genomic data, we have revealed that this recently emerged *L. quercina* population in Colorado is genetically distinct from an established *L. quercina* population in California, suggesting that an accidental introduction of the bacteria from CA to CO is unlikely. Despite clear population differentiation, the results of our analyses were inconclusive whether the CO *L. quercina* population comprises a new taxonomic species. Surprisingly, the CO population had a nucleotide diversity that was twice as high compared to the CA population despite a smaller sample size, suggesting bacterial strains isolated from red oak may be moving to CO from elsewhere.

### Different populations or species?

Despite increased access to molecular and genomic data on various microorganisms, including bacteria, taxonomic placement of species in many cases remains a challenging task (22). Based on previous neighbor-joining phylogenetic analysis of *Lonsdalea* spp. and estimates of average nucleotide identity (ANI) between two previously published genomes of *Lonsdalea quercina* (*L. quercina* ATCC 29281 sampled in CA and *L. quercina* NCCB 100490 sampled in CO (23), Li et al. (24) hypothesized that these two strains from North America may comprise different species (ANI = 94.04%). In this study using population data from both locations, maximum likelihood genealogies based on either 20 or 51 core genes placed both populations within the genus *Lonsdalea*. In the phylogeny, the CA and CO populations formed two distinct and well supported clades that are closely related to *L. iberica*. However, ANI calculated at the population level (all vs all) indicated higher ANI than the one reported by Li et al (24). Our analyses demonstrated that these two populations show genetic separation just above the proposed ANI species boundaries (average ANI = 95.17%, range 94.83 – 95.36%) and cannot be separated into two distinct species based on this measurement. At the same time *L. quercina* from CA and CO show clear signs of differentiation, likely due to a long period of isolation from one another, but they still may be in the process of allopatric speciation. Our pangenome analysis indicated that while CA and CO share most of the core genome (2370 genes), each population has a unique set of core genes (245 core genes in CA and 372 core genes in CO). Some epidemiological differences also exist between the CA and CO populations – *L. quercina* in CA infects acorns of native oaks (*Quercus agrifolia, Q. parvula*, and *Q. wislizeni*), while in CO *L. quercina* infects branches in introduced and planted *Q. rubra* and *Q. palustris*, but acorns in natural areas of native *Q. gambelii*. Overall, these results remain inconclusive regarding the taxonomy of CA and CO populations, and further phenotypic and biochemical assessments are required to clarify their taxonomic placement, therefore in this study we refer to them as populations.

### Clonality and distribution of Lonsdalea quercina populations in California and Colorado

Many plant pathogens spread via clonal reproduction (25, 26), including important bacterial crop pathogens (for example, (27, 28)). Phylogenetic and recombination analyses indicated that both populations of *L. quercina* in this study are comprised of mostly clonal modes of reproduction. For bacteria that rely on external factors for dissemination, the distribution of clones may provide insights into organismal population dynamics. In CA where drippy nut disease caused by *L. quercina* has been established for several decades (first reported in 1967, (14) some clonal isolates were randomly distributed across a large sampled region in CA, including isolates of a genotype B107 that were sampled from trees of *Q. agrifolia* grown in built areas 85km from each other. Genotypic diversity did not differ between isolates from natural and built areas in CA (∼88% in each). There was no association among genotypes with either habitat or host species in CA, suggesting movement of the pathogen between different habitats. These results are not surprising since all oak species in CA are native to the region, and *Q. agrifolia* is a major component of native forests and is also used as a shade tree in residential areas. To date there are no phytosanitary measures established for *L. quercina* in CA, making it possible for the inoculum being moved across large distances by insects, birds, small mammals, and human-mediated activities.

In CO, *L. quercina* of the same genotype were always sampled from planted trees growing in a close proximity to each other (often neighboring trees). Albeit a small population of three, *L. quercina* collected from native *Q. gambelii* were genetically different from isolates collected from planted *Q. rubra*. Whether these differences are due to geographic distance or host species remains unknown. It is known that *L. quercina* from *Q. rubra* is pathogenic on two other introduced to CO species of oaks, *Q. shumardii* and *Q. palustris* (13). In this study, one isolate from *Q. palustris* clustered together with isolates from *Q. gambelii*, suggesting possible movement of bacteria among different host species. Interestingly, the location of sampled *Q. palustris* tree was 45km away from sampled plants of native *Q. gambelii*. Based on population genetic clustering, none of the sampled isolates from *Q. rubra* were genetically similar to the isolates from *Q. gambelii*, and it remains to be determined whether *L. quercina* from native *Q. gambelii* serves as an inoculum reservoir for planted *Q. rubra* hosts or vice versa. Further studies including isolates of *L. quercina* from different host species and geographical areas need to be done to understand the pathogen movement clearer.

Bacterial pathogens can be associated with insects, whereby entering host tissues via feeding sites, wounds, and/or for dissemination among hosts (18, 29–31). It was suggested that in CA, *L. quercina* may enter host tissue via wounds made by acorn weevils, filbertworms, and some cynipid wasps (32). Large numbers of free-living nematodes *Panagrellus redivivus* were often found feeding on ooze produced by *L. populi* on poplar trees in Hungary (18). *Lonsdalea quercina* is also associated with several species of insects in CO that may be helping bacteria spread among host trees (33). Among the associated insects in CO, kermes scale is another main player in drippy blight disease in addition to *L. quercina* (13). This insect does not travel long distances, and this may be one of the explanations why the distribution pattern of recently emerged *L. quercina* genotypes is limited to nearby trees.

### Diversity in a recently emerged population and endophyte hypothesis

Despite more recent emergence, the CO population had ∼2 times higher nucleotide diversity compared to CA population. This is further unexpected because the sampling area in CA was much larger compared to CO (280km and 82km long in CA and CO, respectively and with the majority of CO samples being collected within ∼40 km radius). At the same time, the relative rate of recombination to mutation was the same in both populations. Higher nucleotide diversity observed in CO may suggest the presence of migrants from different source populations. Most isolates in CO were collected from introduced red oak (*Q. rubra*) which is widely planted as a shade tree in the state. Seedlings of red oak are sold in local CO nurseries, but often these seedlings are re-sold from suppliers located in northeastern states of the country where this species is native rather than grown in CO. Several bacterial tree pathogens are known to exist in host tissue without causing symptoms and are therefore spread through grafting or nursery stock [e.g., a bacterial pathogen causing watermark disease in willow trees *Brenneria salicis* (30)]. Other members within the family *Enterobacteriacea, B. gudwini* and *G. quercinecans* associated with acute oak decline, were found in low abundance in healthy trees (34). These two species as well as *L. britannica* and *R. victoriana* were detected in healthy tree tissues using metagenomics (35), suggesting their possible endophytic behavior. There is no published data on *L. quercina* latency in host tissue, but preliminary metagenomics data indicates the presence of *L. quercina* in asymptomatic tissue, suggesting the bacteria may also persist in the host tissue as an endophyte (Raymond & Stewart, unpublished data), and therefore possibly moving with the tree stock. This hypothesis needs to be tested in future studies, that will include identifying *L. quercina* populations from northeastern USA, a native habitat of *Q. rubra*.

### Possible causes of drippy blight outbreak

In CO bacterial isolates were collected from mature oak trees that were planted at least 50 years ago, while first drippy blight diseased trees were noticed in early 2000s. Hence, even if *L. quercina* strains were accidentally introduced with trees to CO, it is unlikely these introductions were the cause of the current epidemic outbreak. In 1990s-2000s three new species of *Lonsdalea* were characterized in different parts of the world. *Lonsdalea britannica* was isolated from symptomatic oak tissues associated with acute oak decline in Great Britain (17), *L. iberica* associated with bark canker of *Quercus* species in Spain (17), and *L. populi*, a causal agent of bark canker of poplar species was reported in Hungary, China, and Spain (18, 19, 36). During the same time symptoms of drippy blight disease were first noticed in Colorado, USA (13). Therefore, it is likely that other environmental factors might have led to the emergence of novel bacterial tree pathogens from the genus *Lonsdalea*.

Bacterial-plant relationships can be altered directly, e.g., via introductions or by changes in environment, or indirectly, e.g., by changes in population dynamics of other organisms with which bacteria interact. According to Filippo et al. (37), in Italy Turkey oak (*Quercus cerris*) growth decline is associated with climate change. Sun et al. (38) reported that warming temperatures and increasing drought significantly affect *Quercus* species richness and distribution in China. Using bioclimatic modeling, Mclaughlin & Zavaleta (39) found that saplings of California valley oak (*Q. lobata*) are especially susceptible to warming and drying conditions (39). Trees stressed by environment conditions can become more susceptible to other stressors, including pathogens (40). It is possible that rapid changes in climate might have altered relationship between *Lonsdalea* spp. and their hosts. This is particularly likely when considering the drippy blight outbreak in Colorado, as these trees are planted outside of the climatic range where they are native.

The kermes scale (*A. galliformis*), now associated with drippy blight in CO, has been present in the state for a long time but was not recognized as a pest because of a lack of significant damage to oak trees. A spike in insect population size was observed with the first drippy blight outbreak (13). Sitz et al (13) hypothesized that interactions between *L. quercina* and *A. galliformis* may be indirect, with the scale possibly acting as a stressor of the host and facilitating growth and spread of bacteria within the host tissue. A similar observation was reported about the wood-boring beetle *Agrilus biguttatus* and *Brenneria gudwini* relationship, in which presence of the beetle larvae triggers the upregulation of tree damaging genes in the bacterium leading to acute oak decline symptoms in Britain (41). Factors leading to the increase of abundance of *A. galliformis* in CO are unknown. It remains to be determined whether the increased scale population and associated increase in the number of entry points through feeding sites has led to the increase of bacterial populations, or vice versa. But it is possible that increase in population size of one organism (for example, caused by climate change) could have caused increase in population size of another organism.

### Conclusions

This study revealed that populations of *Lonsdalea quercina* in California and Colorado are different, and it is unlikely that the drippy blight epidemic outbreak in CO was the result of accidental introduction of the pathogen from CA. Bacteria were isolated from planted stands of introduced *Quercus rubra*, as well as three trees of native *Q. gambelii*, suggesting that the pathogen is also present in natural stands of native CO oak species. However, three isolates from *Q. gambelii* were genetically different from the isolates collected from *Q. rubra*. Due to a small sample size on *Q. gambelii* it remains unclear whether the pathogen moved from native *Q. gambelii* to introduced *Q. rubra* in CO.

While the presence of localized clonality in the CO population is consistent with recent pathogen emergence, higher nucleotide diversity suggests that *L. quercina* isolates from *Q. rubra* might be migrants from elsewhere, possibly from northeastern (NE) states of the USA where red oak seedlings are being produced. Further population genetic analyses, including more populations from *Q. gambelii* in CO and *Q. rubra* in NE USA, are needed to clarify possible causes and sources of the recent outbreak of drippy blight in CO, USA.

## METHODS

### Bacterial strains and DNA extraction

In 2018 oozing acorns and/or acorn caps (California, CA) and oozing branches (Colorado, CO; except for *Quercus gambelii* oozing acorns and acorn caps) were collected from total of 83 (52 CA and 31 CO) trees of various oak species native to North America from trees in undeveloped habitats such as forests, parks, nature preserves, etc. (referred here as “natural areas”) or developed habitats such as residential neighborhoods of cities, parking lots, yards of private houses in rural areas, etc. (referred here as “built areas”) (Table S1). Three symptomatic samples, for both branch and acorn/acorn caps were surface disinfested by dipping in 10% bleach for 1min followed by 1 rinse in distilled H_2_O for 60sec and blotted dried. Branch bark and cambium were removed at canker margin and canker tissue was streaked onto nutrient agar (Sigma-Aldrich, St. Louis, MO, USA) plate. Plates were incubated at 25C for 3-5 days. All cream-colored bacteria were isolated into pure culture on new plates with nutrient agar.

*Lonsdalea* bacterial colonies were verified using colony PCR: bacterial colonies were lysed in 20 μl sterile water at 95°C for 5 min and then used in PCR. Successful amplification of a gryB gene region using species-specific primers [gryB forward: 5’-CTGTACAAGGTGAAGAAAGG-3’, gryB reverse: 5’-CGTCACCAG CATCTCCATCC-3’, (42)] was used in PCR species verification. PCR conditions followed Sitz et al. (33) and were: 94C for 3min, 30 cycles at 94C for 30sec, 60C for 30sec, and 72C for 1min, then finally 72C for 7min using a MJ PTC-200 thermocycler (Bio-Rad Laboratories, Waltham, MA). PCR products were visualized via Sub-Cell GT Wide Mini electrophoresis system (BioRad, Hercules, CA, USA) that yielded a single band (286 kb in length) on 1.5% agarose gel.

Plates with pure cultures of verified *Lonsdalea quercina* isolates were incubated at 25°C for 3-5 days for DNA extraction. DNA was extracted from pure bacterial colonies using Quick-DNA™ Fungal/Bacterial Miniprep Kit (Zymo Research, Irvine, CA, USA) following manufacturer protocol. DNA quality and concentration were estimated with NanoDrop One C spectrophotometer and Qubit 4^MT^ with a dsDNA HS Assay Kit (Invitrogen, Carlsbad, CA, USA). Total of 83 *Lonsdalea quercina* strains (one strain per tree) were subjected to whole genome sequencing. Bacterial cultures were put in 15% glycerol for long term storage at - 80°C.

### Genome sequencing, assembly, and annotation

Illumina short read genome sequencing of 83 isolates of *Lonsdalea quercina* was performed at the Novogene Bioinformatics Institute (Beijing, China). DNA libraries were constructed using NEBNext Ultra DNA Library Prep Kit according to the manufacturer’s protocol. Libraries were sequenced on Illumina NovaSeq 6000 platform with 150 bp paired end reads. Raw data cleaning (adapter removal) and quality control were performed at Novogene.

Raw reads were concatenated, duplicates were removed with BBtools function “dedupe” followed with “reformat” (BBMap – Bushnell B. – (4, 10, 11)). Reads quality was assessed with FASTQC v.0.11.9 (https://www.bioinformatics.babraham.ac.uk/projects/fastqc/). Genome assembly, assembly quality control, and genome annotation were done using Reads2Resistome pipeline v0.0.2 (43). Within the pipeline, genome assembly was performed using Unicycler v0.4.9 (44), genome quality control was assessed using QUAST V5.2.0 (45) and genome annotation was done using Prokka v1.14.5 (46). The pangenome analyses were carried out with Roary pipeline v.3.13.0 (47) with gff3 input files from Prokka annotation. Three datasets of core genes were used for further analyses that are described in detail below (Fig. 5a and b). All core gene datasets were obtained with individual runs of Roary.

**Figure 5.**
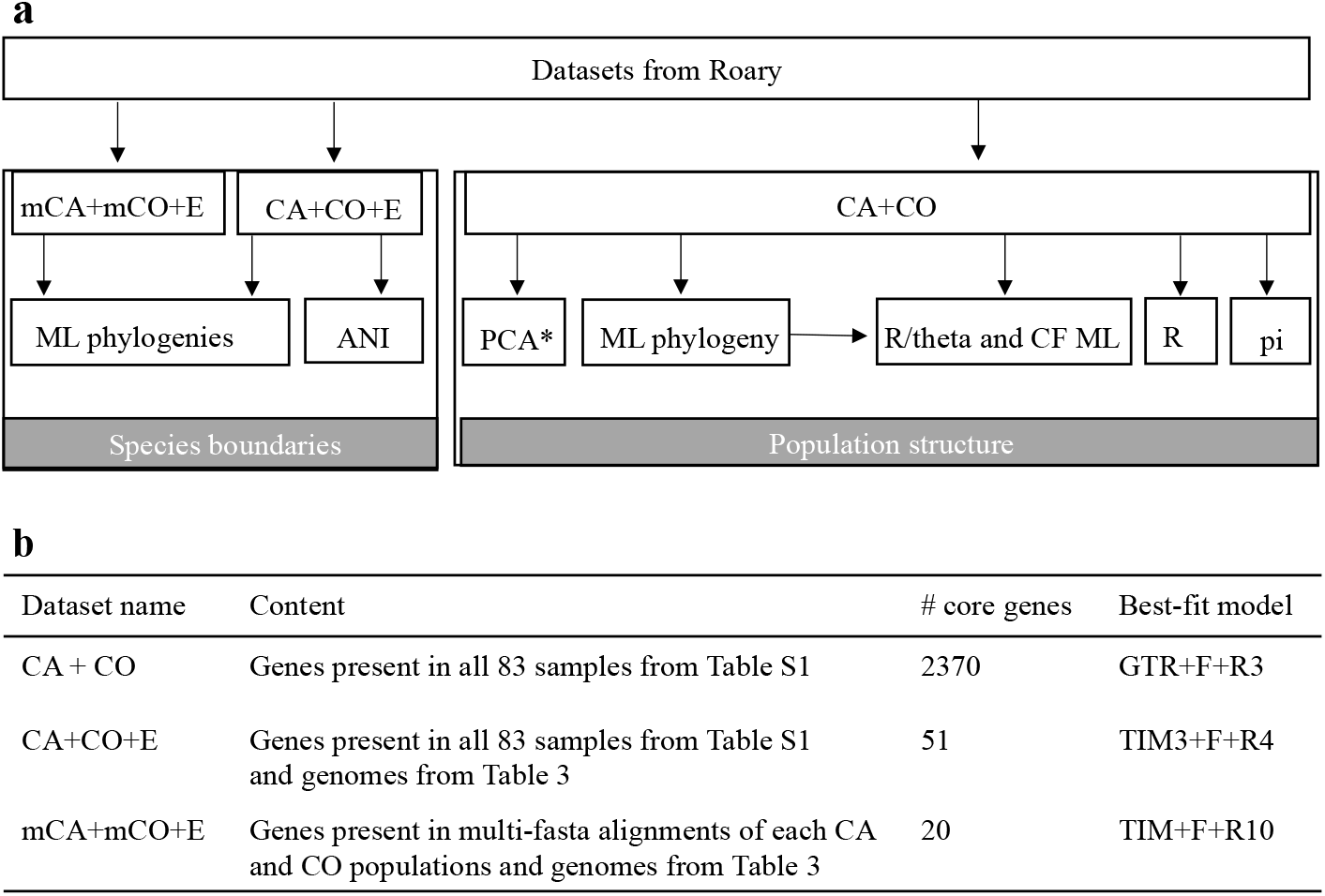
Analyses pipeline using three core genome datasets **(a)** and the datasets description **(b). a**. Arrows indicate what analyses were performed with corresponding datasets. ML phylogeny stands for maximum likelihood phylogenetic analysis with IQTREE v. 2.0.3; ANI – average nucleotide identity estimated with JSpeciesWS; PCA – principal component analysis with Adegenet v. 2.1.3; Recombination – Split Decomposition Analysis for evidence of recombination with SplitTree; R/theta and phylogenetic tree with corrected branch length (CF ML) with ClonalFrameML; pi – unbiased nucleotide diversity estimated with Pixy v.0.95.2. **b**. Datasets were obtained with Roary v.3.13.0. Best-fit model for phylogenetic analyses was selected with ModelFinder. *PCA was conducted on thinned CA+CO dataset (see Materials and Methods for details).

### Species boundaries

The maximum likelihood phylogenetic trees based on core genes were constructed with IQTREE v.2.0.3. (48). First, we determined the phylogenetic position of CA and CO populations within genus *Lonsdalea* and family *Enterobacteriacea*. The genomes of other members of *Enterobacteriacea* family were retrieved from the National Center of Biotechnology Information (Table 3). A total of 51 and 20 core genes present in 100% of samples were used to build phylogenetic trees with all CA and CO samples (CA+CO+E dataset) and with multi-fasta alignments of core genes of each population (mCA+mCO+E dataset), respectively (Fig. 5b). The best-fit model (TIM3+F+R4 – for CA+CO+E dataset, TIM+F+R10 – for mCA+mCO+E dataset, Fig. 5b) selected by ModelFinder (49), was used for the ascertainment bias correction (+ASC) model (50). Tree branch support was assessed via 1000 ultrafast bootstrap replicates (51) with hill-climbing nearest neighbor interchange search bootstrap optimization to reduce the risk of overestimating branch supports (-bnni) due to severe violations of the model. The concatenated trees were visualized with iTOL online tool v.5.7 (itol.embl.de, (52)). Average nucleotide identity (ANI) among all strains was estimated with JSpeciesWS using a BLAST-based approach (ANIb) (53).

**Table 3.**
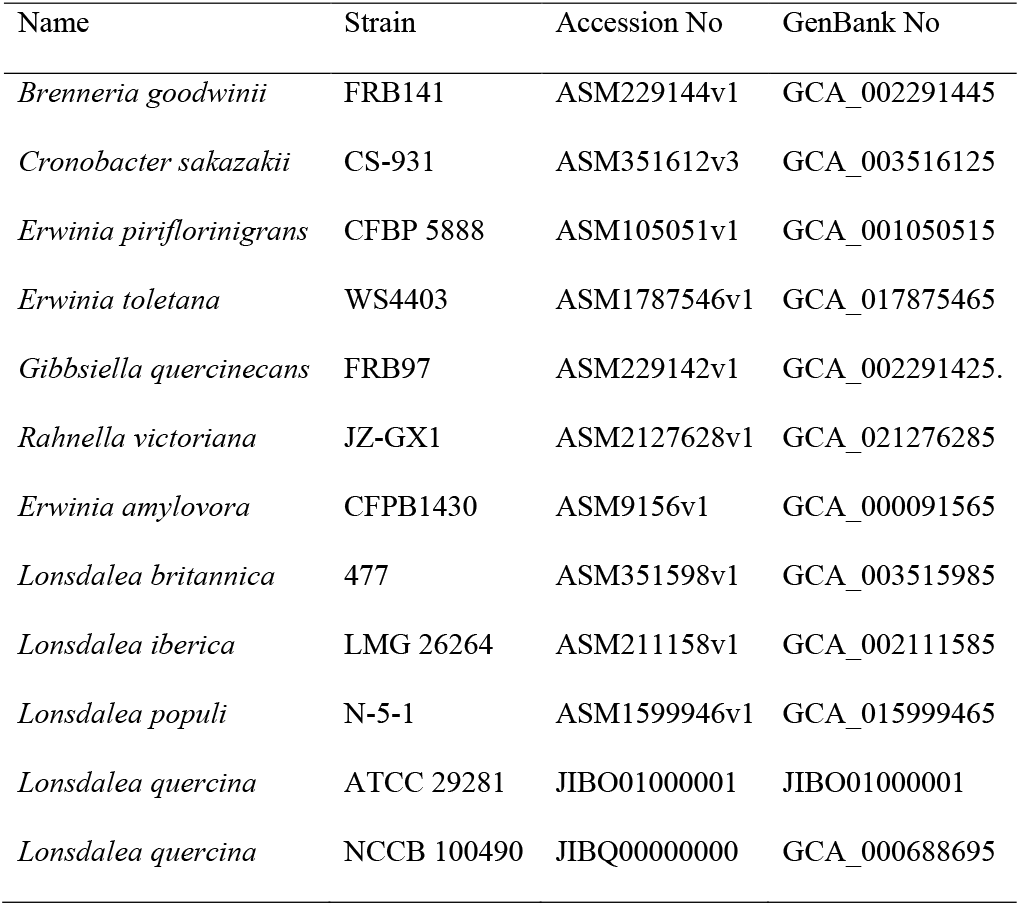
List of genomes of other members of *Enterobacteriacea* family used in this study

### Population structure

Since some of the strains were clonal (see results), further analyses were performed on the clone-corrected dataset (containing 1 representative of each genotype). Population structure was assessed with principal component analysis (PCA) and phylogenetic analysis. PCA was conducted using 2355 unlinked SNPs retrieved from core genes of CA + CO dataset. SNPs were extracted with SNP-sites software (54) followed by filtering with VCFtools v.0.1.16 (55): 1 SNP in every 1000 SNPs (--thin 1000) with minimal frequency 0.05 (--maf 0.05). Analysis was run within R package Adegenet v.2.1.3 (56). An unrooted phylogenetic tree of CA+CO dataset was built to assess population structure within samples obtained in this study. The phylogenetic analysis was conducted as described above, with the best-fit model GTR+F+R3 (Fig. 5b). Unbiased nucleotide diversity (pi) was estimated with Pixy v.0.95.02 (57).

### Recombination

Evidence of recombination was obtained with Split Decomposition analysis and the Phi test implemented in SplitsTree (21). ClonalFrameML (58) was used to estimate relative rates of recombination to mutation (R/theta) in each CA and CO populations and to construct phylogeny with branch lengths corrected to account for recombination. Maximum likelihood phylogenetic tree with CA+CO dataset was used as input for ClonalFrameML.

## Supplemental material

Supplemental materials are available online.

## Data availability

Genome sequence data used in this study have been deposited to the NCBI Sequence Read Archive under project number PRJNA924752 (SAMN32769485 to SAMN32769567). Three datasets used in this study are available on Dryad https://doi.org/10.5061/dryad.m63xsj46h.

## Acknowledgements

Funding was provided by National Park Service (G18AC00335) to JES and RAS. The authors would like to thank everyone who helped locate diseased trees for sample collection. In Colorado: Kathleen Alexander (City of Boulder Forestry), Vince Aquino (University of Colorado), Michael Butterfield (Douglas County Parks and Recreation), Jacque Chomiak (City of Aurora Forestry), Dr. Whitey Cranshaw (Colorado State University), Rob Davis and Richard Wilson (City of Denver Forestry), Thomas Duff (Colorado Parks and Wildlife), Steve Geist and Becky Wegner (SavATree Consulting Group), Carol O’Meara (Colorado State Extension), and Kyle Sylvester (City of Brighton Forestry) as well as in California: Drs. Elizabeth Bernhardt and Ted Swiecki (Phytosphere Research), Phil Cannon (USDA Forest Service), Dr. Walt Koenig (Hastings Natural History Reservation), and Sherby Sanborn (Sherby Sanborn Consulting Arborist). Kristen Otto and Hope Raymond provided laboratory assistance, Hope Raymond and Jill Baty provided field assistance.

JES, RAS, and ISP designed the study, RAS and ISP collected the samples, RAS and LL processed samples, OK analyzed data and wrote the manuscript, RW and JIC contributed to data analyses and data visualization, JES, RAS, ISP, OK, RW, JIC, and ZA contributed to editing the manuscript.

